# De novo Design of A Fusion Protein Tool for GPCR Research

**DOI:** 10.1101/2024.09.14.613090

**Authors:** Kaixuan Gao, Xin Zhang, Jia Nie, Hengyu Meng, Weishe Zhang, Boxue Tian, Xiangyu Liu

**Affiliations:** State Key Laboratory of Membrane Biology, Tsinghua-Peking Center for Life Sciences, School of Pharmaceutical Sciences, Tsinghua University, Beijing, China; Beijing Frontier Research Center for Biological Structure, Beijing Advanced Innovation Center for Structural Biology, Tsinghua University, Beijing, China; Department of Obstetrics, Xiangya Hospital Central South University, Changsha, China

## Abstract

G protein-coupled receptors (GPCRs) play pivotal roles in cellular signaling and represent prominent drug targets. Structural elucidation of GPCRs is crucial for drug discovery efforts. However, structural studies of GPCRs remain challenging, particularly for inactive state structures, which often require extensive protein engineering. Here, we present a de novo design strategy termed “click fusion” for generating fusion proteins to facilitate GPCR structural studies. Our method involves the rational design of structurally stable protein domains rigidly linked to GPCRs. The resulting fusion protein enhances the thermostability of the target GPCR and aids in determining GPCR structures via cryo-electron microscopy (cryo-EM). We further demonstrate that the designed fusion protein can be transferred among structurally similar GPCRs with minor adjustments to the linker region. Our study introduces a promising approach for facilitating GPCR structural studies and advancing drug discovery efforts.

## Introduction

G protein-coupled receptors (GPCRs) constitute a prominent class of membrane proteins distinguished by their seven-transmembrane structure. They serve as key regulators of intracellular biological responses, detecting a wide range of extracellular signals, including hormones, chemokines, light, and neurotransmitters^1^. Given their ubiquitous presence and intricate functional roles within the human body, GPCRs are closely associated with the initiation and progression of numerous diseases^2^. Furthermore, owing to their diverse array of targets and regulatory mechanisms, they emerge as crucial pharmacological targets. Remarkably, over 30% of commercially available drugs rely on modulating GPCR signaling to address a spectrum of ailments such as hypertension and depression^3^.

Among all clinical drugs, GPCR antagonists constitute a large proportion. Presently, a total of 258 GPCR antagonists have garnered approval as clinical drugs, representing 53% of clinically targeted GPCR drugs^3^. Many additional antagonists are currently in clinical trials. For example, atropine serves as an antagonist of muscarinic acetylcholine receptors, playing a pivotal role in various medical applications such as the treatment of bradycardia and as an antidote for specific types of poisoning^4^. PD168368, a representative NMBR antagonist, is considered a promising candidate for inhibiting breast cancer metastasis and invasion^5^. Balovaptan, an inhibitor of 5-HT_2B_R, is clinically used to treat autism spectrum disorder^6^. It is worth emphasizing that determining the structures underlying the interaction between the inhibitors and GPCRs is crucial for understanding signal transduction, physiological mechanisms, and drug design pertaining to GPCRs.

In single-particle cryogenic-electron microscopy (cryo-EM) studies, determining the structures of agonist-bound active-state GPCRs typically involves a 90-150 kDa component comprising the target GPCR and a heterotrimeric G protein. This arrangement provides a stable feature for particle alignment. However, acquiring structural information for antagonist-GPCR complexes is more challenging, particularly for class A GPCRs. These receptors, which constitute the largest family, typically have a smaller molecular weight ranging from 37 to 50 kDa and lack a rigid soluble domain for particle alignment. To address this challenge, a key solution involves inducing a fiducial marker that is not encased by detergent micelles or nanodiscs, such as a nanobody or a fusion protein. Recently, various innovative strategies have emerged to overcome the challenge of projection alignment in inactive-state GPCR structure determination. Building upon a kappa-opioid receptor nanobody, designated as nanobody 6 (Nb6), Michael et al. stabilized the receptor in its inactive conformation. This was achieved by engineering a chimeric receptor capable of interacting with Nb6 through the replacement of the third intracellular loop (ICL3) from a target GPCR with that from the kappa-opioid receptor^7,8^. However, not all chimeras can be readily expressed and purified, posing challenges for their utilization. In X-ray crystallography studies, the T4 lysozyme (T4L) fusion strategy has emerged as a robust method, widely used for successfully determining the structures of inactive-state GPCRs^9,10^. However, this approach is less effective in cryo-EM studies, likely due to the small size of T4L and its flexibility relative to TM5/6. To address these limitations, various fusion strategies have been designed and implemented, yielding promising results in the cryo-EM field^8,11-13^. These methods emphasized the importance of establishing a rigid attachment between the GPCR and the fusion protein. But achieving such a rigid connection is far from trivial and often requires extensive engineering efforts to optimize the construct. Consequently, there is an urgent need to develop a more robust and less labor-intensive method.

With the advent of machine learning, advancements in protein structure prediction and protein design tools have introduced new approaches to address the challenge of rigid fusion protein integration. In this study, we employ the predicted local-distance difference test (pLDDT) from AlphaFold2 as a metric to assess the rigidity of the linker between GPCRs and the fusion protein. Leveraging ProteinMPNN and RFdiffusion^14-16^, we designed de novo fusion proteins for muscarinic acetylcholine 1 receptor (M1R) and neuromedin B receptor (NMBR), achieving atomic-level resolution structures. Additionally, we successfully migrated the M1R fusion protein to the 5-hydroxytryptamine 2B receptor (5-HT_2B_R), thereby overcoming the lack of an inactive state structure. Based on these findings, we introduce a de novo designed fusion protein tool applicable for GPCRs, termed “click fusion”, which may greatly facilitate structural studies for GPCRs through its simplicity, speed, and reliance on in silico experiments rather than extensive experimental screening.

## Results

### Stepwise Generation of Rigid Fusion Proteins for GPCRs

The fusion protein design was achieved through backbone generation followed by side chain packing, as previously reported^16^. The backbone of click fusion proteins (Clip) was generated utilizing the RFdiffusion method^15^. Typically, the AlphaFold model of the target GPCR served as a substructure^17^, with the click fusion protein formed between transmembrane helices 5 and 6 (TM5/6). Initial trials to generate a ∼150 aa fusion protein in a single-step RFdiffusion often resulted in a skeleton that exhibited a tendency to fold towards the transmembrane regions of the GPCRs. To mitigate the unwanted extended scaffold towards the TM region, we implemented a strategy known as stepwise generation. This process involved three steps: first, TM5/6 extension; second, generation of a stabilizing intermediate helix for the elongated TM5/6; and finally, generation of the entire backbone of the click fusion protein. The resulting backbone was then employed for sequence design (Fig. 1).

**Fig. 1.**
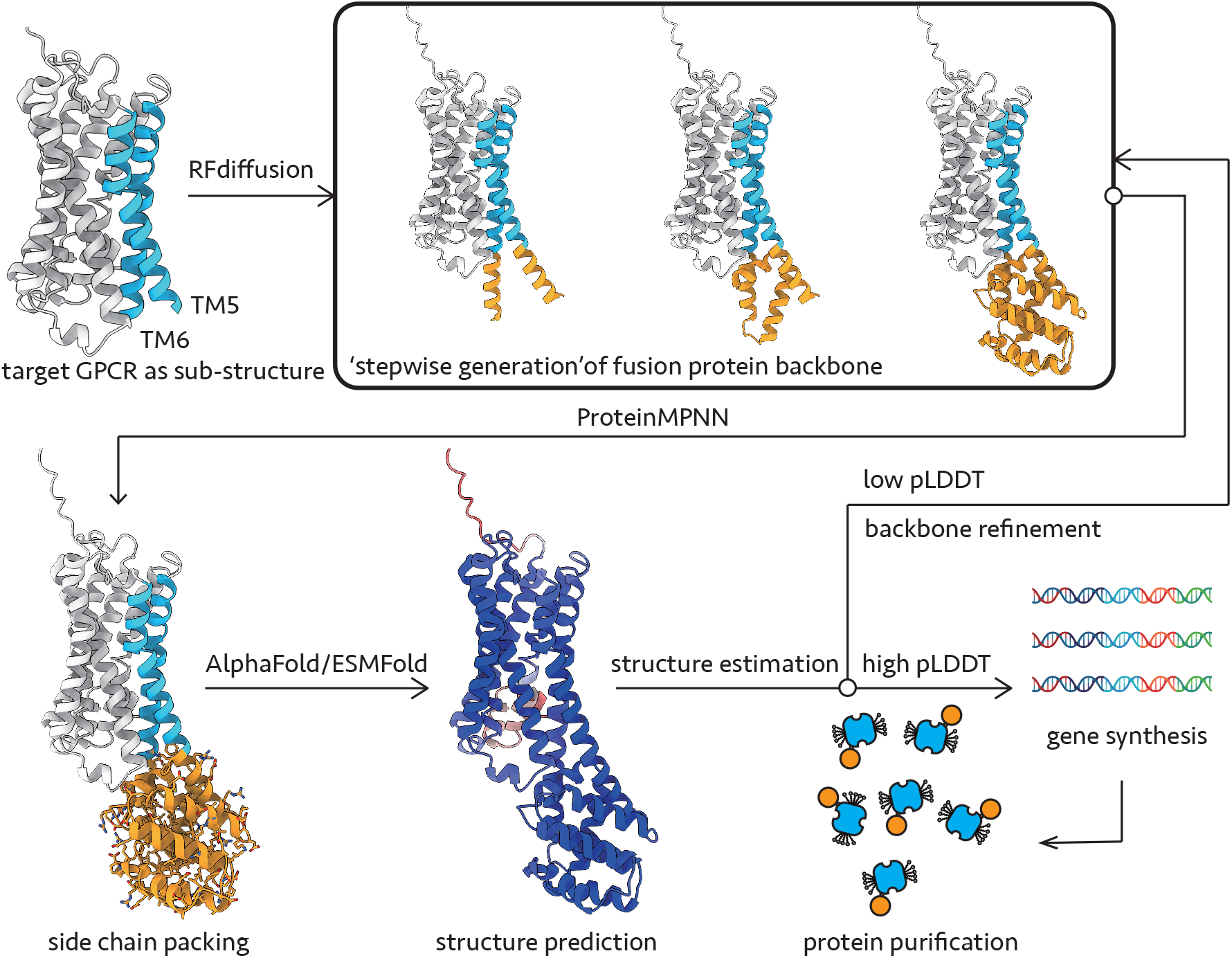
Computational design of click fusion protein (Clip).

Side chain packing of the click fusion protein was performed using ProteinMPNN^16^, followed by structure prediction using AlphaFold2 or ESMfold^14^. The predicted structures were validated using pLDDT to assess sequence rationality and linker rigidity. Structures with lower scores underwent refinement using the RFdiffusion module, while sequences with high confidence were synthesized, expressed, and purified to obtain GPCR-Clip proteins (Fig. 1). Following this process, we initially designed two Clips for two class A GPCRs, M1R and NMBR, with molecular weights of 17 kDa (Clip1, for M1R) and 19 kDa (Clip2, for NMBR), respectively. Notably, both constructs predominantly comprised alpha-helices. These two constructs, M1R-Clip1 and NMBR-Clip2, were then utilized for subsequent experiments.

### Biochemical Characterization of the M1R-Clip1 Construct

M1R is associated with learning, memory, and cognition, making it an attractive target for the treatment of various central nervous system disorders, including Alzheimer’s disease, schizophrenia, and drug addiction^18^. The crystal structures of M1R-antagonist complex were reported in 2016 and 2020: one in complex with tiotropium and the other in complex with muscarinic toxin-7 and atropine^19,20^.

Cell surface staining was conducted to assess the expression levels of M1R-Clip1 and M1R-ΔICL3, an intracellular loop 3 (ICL3) truncated construct with previously demonstrated excellent expression level. Previous studies suggest that M1R-ΔICL3 has higher expression levels than wild type M1R (M1R-wt), while the pharmacological characteristics are similar to M1R-wt^20^. Cell surface staining results indicate that M1R-Clip1 exhibits a similar expression level to M1R-ΔICL3 (Extended Data Fig. 1), suggesting the preliminary feasibility of fusing de novo designed proteins with GPCRs.

To characterize the functional properties of M1R-Clip1, we compared the M1R-Clip1 construct with the wild-type M1R or M1R-ΔICL3. A radioligand saturation binding assay was performed to measure the Kd of [^3^H]-N-methylscopolamine (NMS) to these constructs. M1R-Clip1 exhibits a similar Kd value (0.5 nM) to that of the M1R-ΔICL3 (1.1 nM), indicating that they have comparable affinities for the hot ligand (Fig. 2a, b). We then measured the affinities of a representative agonist and antagonist towards M1R-Clip1 or M1R-ΔICL3 using a radioligand competition binding assay. Interestingly, M1R-Clip1 exhibits higher affinity for the antagonist atropine and lower affinity for the agonist iperoxo than M1R-ΔICL3 (Fig. 2g, h), suggesting M1R-Clip1 is stabilized in an inactive state. Previous structural studies suggest that the outward displacement of TM6 is a hallmark of GPCR activation^21^. However, in M1R-Clip1, the presence of the fusion protein and the intermediate helix prevents TM6 from swinging outward, thereby locking the receptor in an inactive state. This is consistent with isotope binding experiments.

**Fig. 2.**
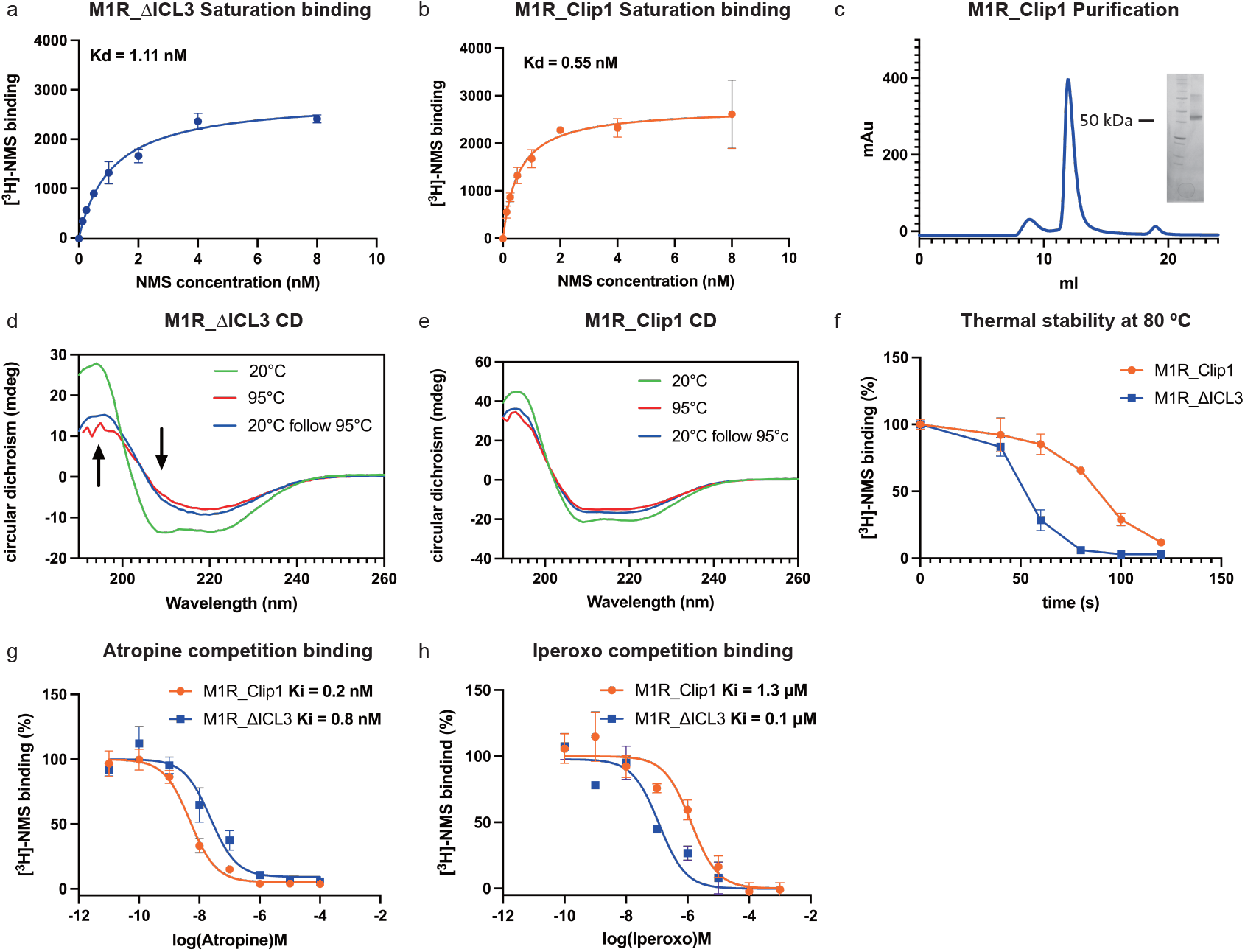
Biochemical Characterization of the M1R-Clip1 Construct. **a, b** [^3^H]-NMS Saturation binding assay for the M1R constructs: M1R_ΔICL3 (**a**) and M1R_Clip1 (**b**). **c** Size exclusion chromatography (SEC) analysis of the M1R_Clip1. The peak fraction was analyzed by SDS-PAGE (insert). **d, e** Circular dichroism spectra of the M1R constructs, M1R_ΔICL3 (**d**) and M1R_Clip1 (**e**) at different temperatures (green,25 °C; red, 95 °C; blue, 95 °C followed by 25 °C). **f** [^3^H]-NMS binding assay at 80ºC of different M1R constructs (blue, M1R_ΔICL3; orange, M1R_Clip1) **g, h** [^3^H]-NMS competition binding assay for different M1R constructs (blue, M1R_ΔICL3; orange, M1R_Clip1) using representative antagonist atropine (**g**) or agonist Iperoxo (**h**). Values represent the means ± SD of 3 independent experiments.

To investigate whether the Clip1 fusion affects the stability of M1R, we assessed the protein’s thermal stability using circular dichroism (CD) spectroscopy and a radioligand binding assay with purified receptors. Both M1R-Clip1 and M1R-ΔICL3 were expressed and purified in a similar way (Methods, Figure 2c, Extended Data Fig. 2g). In the CD spectrum, both M1R-ΔICL3 and M1R-Clip1 exhibit helical signals, consistent with the characteristics of GPCRs and the design of fusion protein (Fig. 2d, e). We then measured the melting temperature (Tm) of these two constructs by recording the spectra at different temperatures^22^. The results show that the Tm of M1R-Clip1 is around 10°C higher than M1R-ΔICL3 (Extended Data Fig. 2c, d).

Notably, the CD spectra reveal that M1R-Clip1 maintains a stable secondary structure even after heating, whereas M1R-ΔICL3 significantly loses its helical structure upon heating, and the secondary structure does not recover with temperature restoration, indicating protein denaturation (Fig. 2d, e; Extended Data Fig. 2a, b). We further verified the receptor’s stability using radioligand binding assays. Upon heating at 80°C, M1R-ΔICL3 rapidly lost its activity, whereas M1R-Clip1 was able to maintain its activity for a longer time (Fig. 2f; Extended Data Fig. 2f). This observation further supports the stability and structural integrity of M1R-Clip1, highlighting its potential utility in further studies.

### Structure Determination of M1R-Clip1

Cryo-EM samples were prepared using the purified M1R-Clip1 protein. Clear features of Clip1 were observed on 2D class averages, even with a small data collection comprising approximately 400 micrographs (Fig. 3a). Utilizing a dataset containing approximately 2,000 micrographs, we achieved a reconstruction with a global resolution of 3.3 Å (Fig. 3b). Notably, the local resolution of the reconstruction is higher in the transmembrane domain than in the soluble domain, as observed in the Nb6 strategy (Extended Data Fig. 3)^8^. Despite the relatively lower resolution of the soluble part, the secondary structure remains clearly distinguishable. Importantly, the experimentally determined structure closely matches the AlphaFold2 predicted structure (Fig. 3c), thereby validating the stability and reliability of the de novo designed Clip. Comparison with the crystal structure indicates that the structures obtained by X-ray crystallography or cryo-EM exhibit no significant differences, with the ligand’s pose closely resembling the shape of the density (Fig. 3d, e, f). In summary, our results demonstrate the feasibility of obtaining high-resolution inactive GPCR structures using de novo designed Clip strategy.

**Fig. 3.**
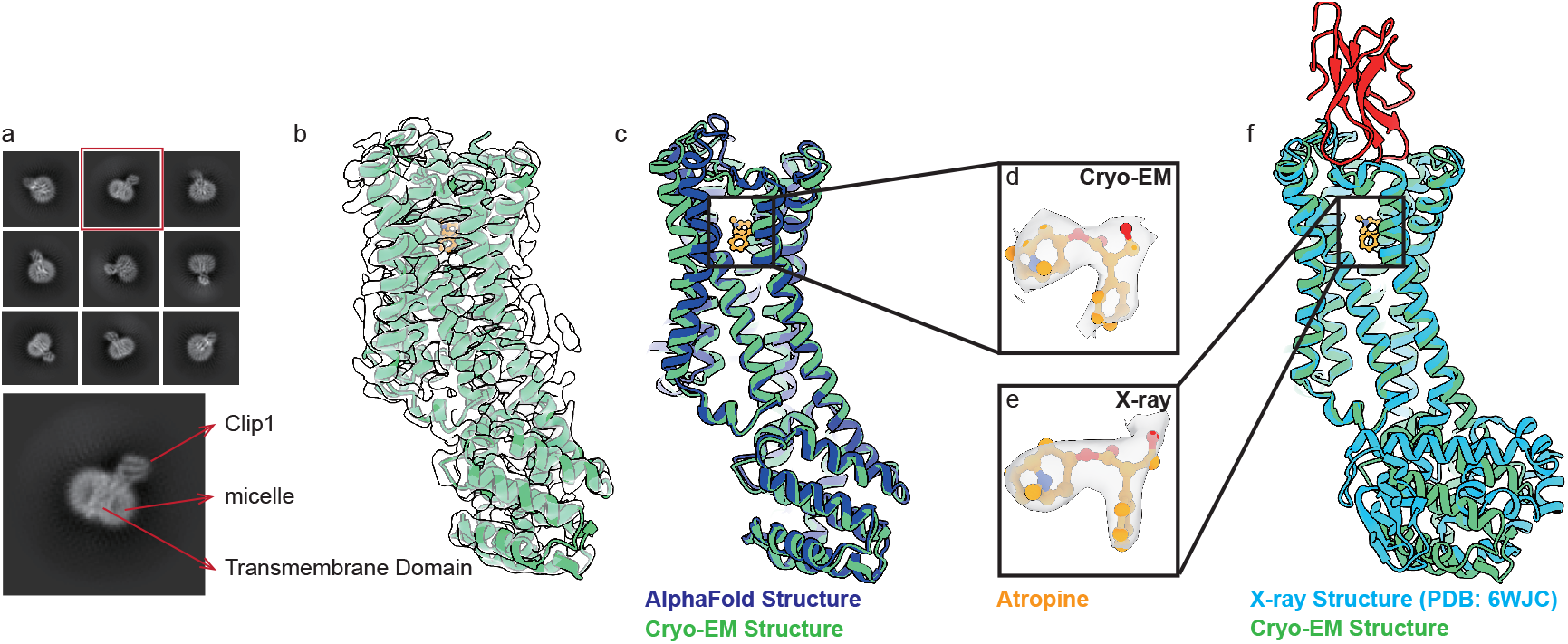
Overall structure of M1R_Clip1. **a** 2D average of M1R_Clip1. **b** cryo-EM structure of M1R-Clip1. **c, f** structure comparison between cryo-EM structure and Alphafold structure (**c**) or X-ray structure (**f**). **d, e** the binding pose and density of ligands from cryo-EM structure (**d**) and X-ray structure (**e**).

### NMBR Structure

To verify the universality of this strategy, we attempted to obtain a structure of NMBR, which has long lacked an inactive state structure. NMBR mediates biological processes in the central nervous system and other tissues through its interaction with neuromedin B^23,24^. Based on the NMBR-Clip2 construct, we determined the inactive-state structure of NMBR in complex with its antagonist PD168368 with a global resolution of 3.4 Å (Fig. 4a).

**Fig. 4.**
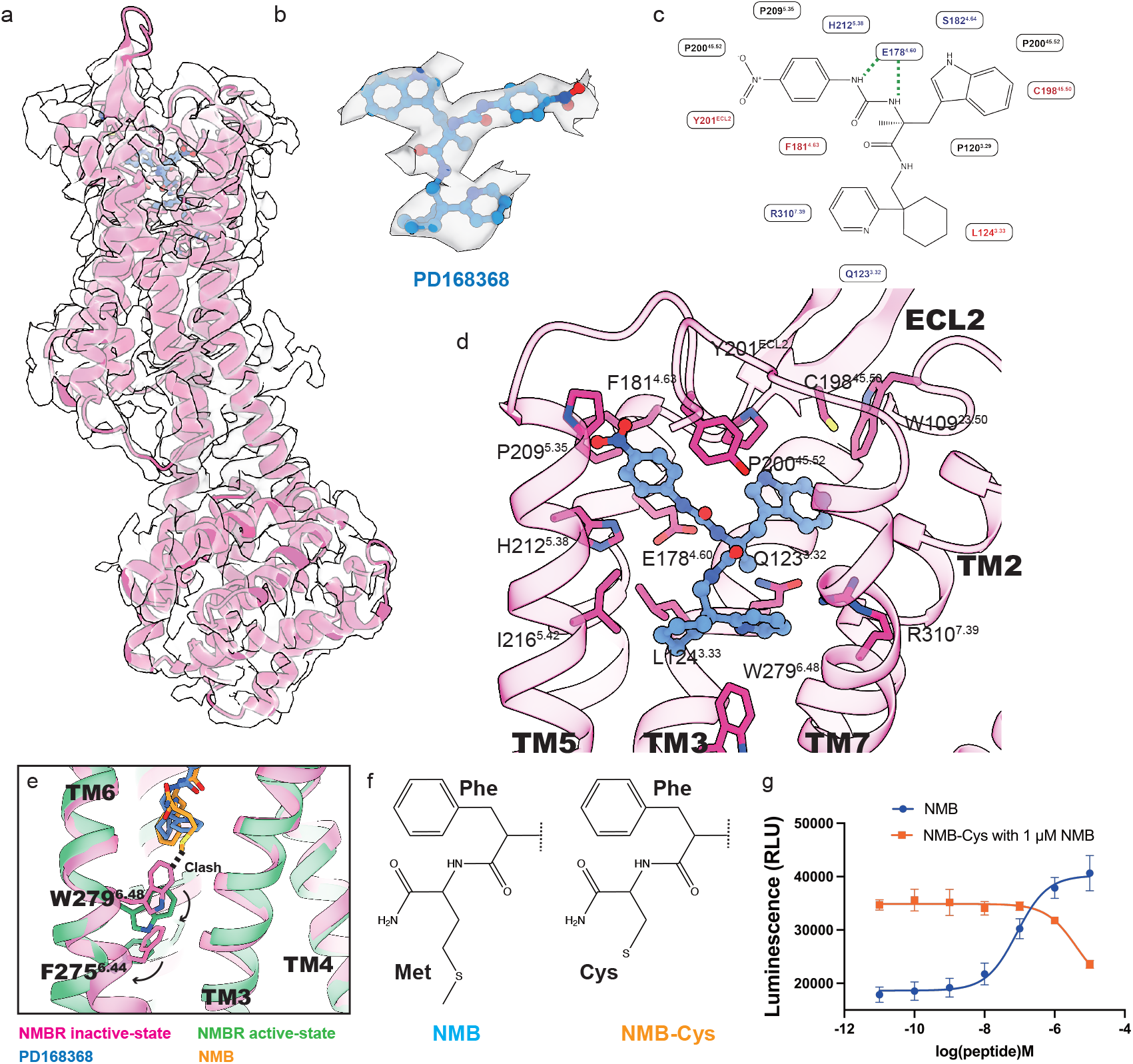
Overall structure and activation mechanism of NMBR. **a** Overall cryo-EM structure of NMBR_Clip2. **b** Density and binding pose of PD168368. **c** Two-dimensional schematic depiction of the PD168368 binding pocket on the NMBR. **d** PD168368 binding pocket. **e** Structure comparison between different states of NMBR (pink, inactive-state NMBR; green, active-state NMBR; blue, PD168368; orange, NMB). **f** Design of NMB-Cys as a NMB analog. The C terminal of NMB (Blue) or NMB-Cys (Orange) are different. **g** NanoBit assay of different ligands (NMB, blue; NMB-Cys with 1 µM NMB, orange) on the NMBR. Values represent the means ± SD of 3 independent experiments.

The structure of Clip2 aligns well with the predicted structure (Extended Data Fig. 4a), further supporting the credibility of our protein design strategy. The orthosteric pocket of the receptor is well resolved by cryo-EM density, with a cloverleaf-shaped density contributed by the antagonist PD168368 (Fig. 4b). We docked PD168368 within this density, and the compound forms extensive interactions with the amino acids of the receptor (Fig. 4c, d). Comparing the structure with the active state NMBR structure (PDB: 8HP0)^24^, we found that both the endogenous agonist NMB and antagonist PD168368 bind to a similar orthosteric pocket. However, since NMB is an endogenous peptide ligand, it binds in a broad pocket, with several residues on its N-terminal side forming extensive interactions with the ECL2 of NMBR. In contrast, the small molecule PD168368 binds in the lower part of this broad pocket (Extended Data Fig. 4b).

Comparison of the active state and inactive state NMBR structures reveals the activation mechanism of the receptor. NMB deeply inserts into the orthosteric pocket of NMBR, with its terminal methionine slightly clashing with the toggle switch W^6.48^ in the inactive receptor. This clash directly drives the downward movement of W^6.48^, transitioning the receptor from inactive to active state. In contrast, PD168368, which binds more shallowly, acts as an antagonist of NMBR (Fig. 4e). To validate this hypothesis, we synthesized a mutant peptide of NMB, replacing the carboxyl-terminal methionine with cysteine. This resulted in an NMB-Cys peptide with a smaller residue than NMB at the carboxyl terminus, which may lead to a shallower binding pose and an inability to shift the toggle switch W^6.48^ (Fig. 4f). Activity assays confirmed our expectations: NMB-Cys cannot activate NMBR and instead exhibits antagonist activity (Fig. 4g). These results further highlight the importance of structural information in guiding the development of novel ligands for NMBR and other GPCRs.

### ‘Click Fusion’ Enabled Structure Determination of Inactive State 5-HT_2B_R

While the de novo design of click fusion proteins represents a relatively simple and rapid process compared to traditional experimental screening, we tested whether it’s possible to transfer Clip from one GPCR to another to offer a plug-and-play approach that minimizes the need for extensive protein engineering. This is inspired by previous work on T4 lysozyme (T4L) fusion constructs, which exhibit strong portability among different GPCRs^9,25,26^. We applied this strategy to 5-HT_2B_R, a GPCR involved in the regulation of the fibrosis disorders, cardiovascular system, cancer, the nervous system and the gastrointestinal (GI) tract^27^. Initially, we transferred Clip1 from M1R to 5-HT_2B_R. These two receptors share a highly similar backbone, particularly in their TM5/6 region. Through structural comparison, we appropriately truncated 5-HT_2B_R, fused the truncated receptor with Clip1, and predicted the structure using AlphaFold2. The predicted structure exhibited high confidence as revealed by high pLDDT values (Extended Data Fig. 6a). In this process, the complementary receptors and Clip act like matching clasps, seamlessly “clicking” together, which underscores the concept of click fusion (Fig. 5a). While the transfer of Clip1 from M1R to 5-HT_2B_R showed promising compatibility, the same cannot be said for Clip2. The poor compatibility observed between NMBR and 5-HT_2B_R suggests that the migration of Clip2 is not flawless, highlighting the need to design different Clips tailored to specific TM5/6 configurations (Extended Data Fig. 6b). The expression level of 5-HT_2B_R-Clip1, detected by cell surface staining as previously described, was similar to that of M1R-Clip1, indicating the feasibility of the plug-and-play approach (Extended Data Fig. 1).

**Fig. 5.**
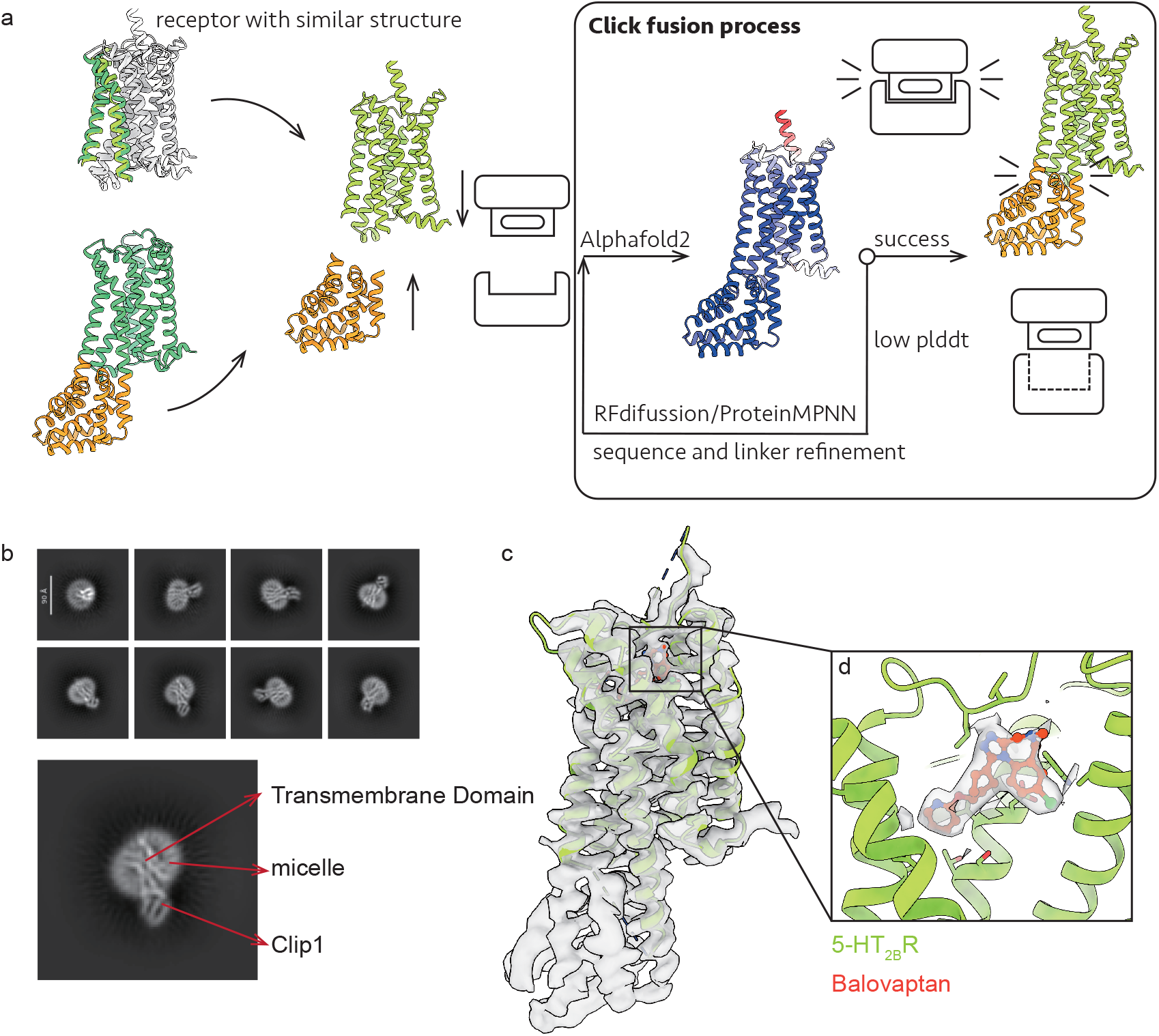
Click fusion strategy and the overall structure of 5-HT_2B_R. **a** The workflow of click fusion strategy. **b** 2D average, overall structure of 5-HT_2B_R and ligand density of balovaptan.

Subsequently, we purified and solved the structure of the 5-HT_2B_R bound with balovaptan, an investigational drug in Phase 3 trials that previously identified as a 5-HT_2B_R antagonist. In the 2D classification of the 5-HT_2B_R-Clip1, we clearly observed the features of the fusion protein, exhibiting well-defined secondary structures (Fig. 5b). The consistency with the characteristics shown by M1R-Clip1 indicates that the Clip fusion strategy is effective. Ultimately, we obtained a 3.6 Å cryo-EM structure for 5-HT_2B_R-Clip1 (Fig. 5c). The binding pose of balovaptan is well-resolved by a density resembling the shape of balovaptan in the orthosteric pocket of the receptor (Fig. 5d). Taken together, the results suggest the Clip fusion strategy could be used for structural determination of inactive state GPCRs and to investigate the interaction between GPCRs and their antagonist drugs.

## Discussion

In this study, we present inactive-state structures of three Class A GPCRs using a de novo design fusion protein strategy, termed “click fusion.” This term encapsulates two main concepts: firstly, protein engineering is predominantly carried out in silico, requiring only the click of a mouse rather than extensive experimental screening; secondly, the transfer of Clips is seamless, resembling the matching of clasps clicking together.

We initially applied this method to M1R, which already had crystal structures available. The quality of the overall structure and ligand density of M1R-Clip1 is comparable to the previously reported X-ray structure. Subsequently, we extended the approach to NMBR, which lacked an inactive-state structure. The successful determination of the NMBR-Clip2 structure highlights the universal applicability of the Clip method. Furthermore, we demonstrated that the Clip fusion protein could be transferred between GPCRs by solving the 5-HT_2B_R-Clip1 structure, with Clip1 directly transferred from the M1R-Clip1 construct. Although the transfer of Clip1 from M1R to 5-HT_2B_R showed promising compatibility, the same could not be applied to Clip2 from NMBR to 5-HT_2B_R. The observed poor compatibility between NMBR and 5-HT_2B_R indicates that the migration of Clip2 is not flawless, underscoring the need to design distinct Clips tailored to specific TM5/6 configurations.

GPCRs are targets for approximately 34% of FDA-approved drugs, with more than half of these being GPCR antagonists. Despite various strategies for determining inactive-state GPCR structures, success rates remain limited, often requiring extensive protein engineering efforts. The click fusion strategy offers a novel approach to generating constructs suitable for inactive-state structure determination, which could greatly facilitate structure-guided antagonist drug development for GPCRs.

Recent advances in drug screening methods have shown promising results for GPCRs, including structure-based virtual screening techniques such as molecular docking^28^ and experimental screening using purified proteins like DNA-encoded libraries^29^. One challenge with experimental screening is obtaining stable purified receptors, as GPCRs are notoriously difficult to work with and their purified forms are often unstable, particularly in the absence of high-affinity ligands. Our data suggest that the Clip-fused M1R exhibits higher thermal stability compared to its wild-type counterpart in the apo form. Additionally, the Clip-fused receptors appear to be locked in the inactive state. These observations strongly suggest that GPCR-Clip constructs are well-suited for hit screening techniques, such as DNA-encoded library screening^29^, or other library display techniques like yeast surface display, phage display, or ribosome display^30-32^. This method may facilitate the screening of novel small molecules, cyclopeptides, nanobodies, or toxins that specifically recognize the inactive state of target GPCRs, offering promising avenues for further research in this field. Furthermore, this method could be readily applied to other small membrane proteins, such as transporters, to aid in cryo-EM structure determination or to generate more stable fusion proteins.

## Methods

### Computational design

The de novo design of Clips was initiated through a collaborative approach integrating RFdiffusion^15^, ProteinMPNN^16^, and AlphaFold2^14^. RFdiffusion facilitated the backbone design utilizing a stepwise generation strategy, using the AlphaFold2-predicted structure of the target GPCR from GPCRdb^3^ as a substructure template. The ICL3 was deleted, and the residues connecting ICL3 and TM5/TM6 were defined as the fusion sites for further substructure generation. Subsequent steps involved generating multiple backbones comprising approximately 10 to 20 amino acids between the fusion sites. The backbones with extended helical conformations of TM5/6 were chosen for next steps, which involved the generation of approximately 20 amino acids to form a stabilizing helix clamped by TM5/6. After that, 100-150 amino acids were generated to complete the Clip backbone. Side chain packing was executed using ProteinMPNN, pretrained soluble model weights was used in this process^16^. AlphaFold2 was then employed for structure prediction. Sequences with high pLDDT scores were kept and regions with low scores were subjected to RFdiffusion for further backbone refinement. Finally, the DNA sequences encoding the designed Clips were synthesized for further studies. For M1R-Clip1, two protein sequences with the same backbone were generated using ProteinMPNN. Both sequences were synthesized and tested for protein expression. Both of the designed protein expressed well and one of them were chosen for structural determination. For NMBR-Clip2, only one protein sequence was chosen for gene synthesis and tested for expression, which exhibited good expression level and was chosen for structural studies.

### Expression and purification of the receptor

The M1 muscarinic receptors, including both M1R-ΔICL3 and M1R-Clip1, NMBR-Clip2 and 5-HT_2B_R-Clip1 (Supplementary Table 2) were cloned into pFastBac vector. Expression of the constructs was achieved using the Bac-to-Bac Baculovirus Expression System (Invitrogen) in Sf9 cells. The cells were infected with baculovirus at a density of 4.0 × 10^6^ to 5.0 × 10^6^ cells per milliliter and harvested after 48 hours of infection. To enhance protein expression, M1R-ΔICL3 was also treated with 10 μM atropine. The purification of both proteins was carried out using Ni-NTA chromatography, Flag affinity chromatography, and size-exclusion chromatography as previously reported^19^.

### Cryo-EM grid preparation

For the preparation of cryo-EM grids, 4 µl of the purified GPCR-Clips described above were deposited onto glow-discharged Au Quantifoil grids. Subsequently, the grids were gently blotted with Whatman No. 1 qualitative filter paper within a Vitrobot Mark IV (Thermo Fisher) maintained at 8 °C and 100% humidity for 4 s, employing a blot force of four, before immersion into liquid ethane.

### Data collection and processing

Cryo-EM data of M1R-Clip1, NMBR-Clip2, and 5-HT_2B_R-Clip1 were collected at Titan Krios G3i TEM (ThermoFisher), acquired with AutoEMation^33^. The data processing was carried out following a similar process using cryoSPARC (v4.5.1)^34^. First, several rounds of 2D classification were conducted. The best 2D averages were selected for 3D reconstruction and were used as templates for picking particles across the entire dataset. A 3D volume with clear soluble features and a well-defined transmembrane domain was selected as the template for further 3D classification. After performing multiple rounds of heterogeneous refinement and non-uniform refinement, a 3D volume with well-resolved secondary structures was obtained. The detergent micelles in the 3D volume were then removed to create a mask that encapsulates the transmembrane domain, which was used for focused classification. After several iterations of heterogeneous refinement and 3D classification, a high-resolution volume was obtained. The dataset for M1R_Clip1 contained 3,884,205 particles, with 121,129 particles used for final reconstruction of the 3.3 Å structure. The dataset for NMBR_Clip2 contained 2,721,525 particles, with 149,635 particles used in the final reconstruction of the 3.4 Å structure. The dataset for 5-HT_2B_R-Clip1 contained 6,384,931 particles, with 291,656 particles used in the final reconstruction of the 3.6 Å structure.

### Model building and refinement

Alphafold2 predicted models of M1R-Clip1, NMBR-Clip2, and 5-HT_2B_R-Clip1 were used as initial structure models. Phenix.elbow (1.20.1-4487) was used to generate the coordinates and chemical constraints for atropine, PD168368, and balovaptan^35^. The models were fitted into the cryo-EM map using UCSF ChimeraX-1.3^36^, followed by manual adjustments using COOT-0.9.8.7^37^. The refinement and validation of the structures were conducted through PHENIX^38^.

### Cell surface staining

Cell surface staining was employed to assess the expression levels of the aforementioned receptors. Specifically, the cells were suspended in phosphate buffer solution (PBS, pH=7.4) supplemented with 2 mM CaCl2 and incubated with Alexa-488 conjugated anti-Flag antibody (diluted in HBSS at a ratio of 1:300, Thermo Fisher, Cat # MA1-142-A488) in darkness for 15 minutes at room temperature. After incubation, the cells were washed twice with PBS prior to fluorescence measurement via flow cytometry BD Accuri C6 (BD Biosciences), with excitation at 488 nm and emission at 519 nm. A standard gating strategy based on cell size and granularity was implemented.

### Radioligand binding assay

The human M1R constructs, including M1R_wt, M1R_ΔICL3, and M1R_Clip, were expressed in Sf9 insect cells or COS7 cells. For membrane preparation, 50 mL of cells were harvested and homogenized in 8 mL of lysis buffer containing 20 mM Tris (pH 7.5) and 1 mM EDTA. The lysate was firstly centrifuged at 800 rpm for 10 minutes, after which the supernatant was collected and subjected to further centrifugation at 18,000 rpm for 20 minutes. The resulting membrane pellet was resuspended in a binding buffer composed of 20 mM HEPES (pH 7.5) and 100 mM NaCl.

To perform the saturation binding of M1R constructs, the membranes were incubated with varying concentrations of [^3^H]-NMS for 1 hour at room temperature in a buffer containing 20 mM HEPES (pH 7.5), 100 mM NaCl, 5 mM MgCl2, and 0.1% BSA. Non-specific binding was assessed using samples containing 10 μM atropine. For the competition binding assay, the affinity of M1R constructs for atropine or iperoxo was measured by incubating the membranes with a fixed concentration of [^3^H]-NMS and varying concentrations of ligands for 1 hour at room temperature. For the thermal stability assay, the purified M1R_ΔICL3 and M1R_Clip1 proteins were reconstituted into high-density lipoprotein (HDL) particles, which were composed of apolipoprotein A1 and a lipid mixture of POPG and POPC in a 3:2 molar ratio. The reconstituted proteins were then subjected to heat treatment at 80°C for different time, followed by incubation with [^3^H]-NMS for 1 hour. Reactions were harvested using rapid filtration through GF/B filters using Brandel 48-well harvester, and the data were analyzed with Prism7^25,39^.

### Circular dichroism

Circular dichroism (CD) experiments were performed using a Chirascan plus (Applied Photophysics Ltd) equipped with a multi-cell holder that allowed for precise temperature control. Wavelength scans were initially taken from 260 to 190 nm at 25°C and 95°C, followed by additional scans at 25°C after a cooling process, which took around 5 minutes. For the temperature melting experiments, the CD signal at 222 nm was monitored, with the temperature increased by 1°C per minute and a 30-second equilibration period at each step. These measurements were conducted using a protein concentration of 0.2 mg/mL in HEPES/NaCl buffer (20 mM HEPES, 100 mM NaCl, pH 7.4), employing a 1 mm path-length cuvette. The melting temperatures were extracted by fitting the collected data to a sigmoidal curve.

### NanoBit G protein association assays

NanoBit G protein association assays were performed as previously described using plasmids provided by Prof. Asuka Inoue^40^. For measuring Gq signaling, HEK293T cells were transfected with a plasmid mixture containing 500–1500 ng LgBiT-inserted mini-Gsq subunit and 500–1500 ng NMBR-SmBiT receptor^41^. After 24 hours, luminescence was measured following ligand addition. For the inhibitor assay, cells were stimulated with 1 µM NMB after incubation with NMB-Cys. Data were analyzed using Prism 8, and statistical analysis was conducted using a t-test.

## Supporting information

Extend data

## Author information

### Contribution

K.G. developed the computational method and design the fusion protein with help from X.Z.; K.G., X.Z., J.N. and H.M. preformed protein expression and purification; X.Z. and K.G. collected the cryo-EM data and structure determination and refinement; K.G. and X.Z. characterized the biochemical properties of M1R; J.N. characterized the biochemical properties of NMBR; X.L. coordinated the experiments. X.L., B.T. and S.Z. oversaw the overall study; The paper was written by X.L. and K.G. with input from J.N. and Z.X; All authors contributed to the editing of the paper.

