## Supplementary material for "De novo Design of A Fusion Protein Tool for GPCR Research": Extend data

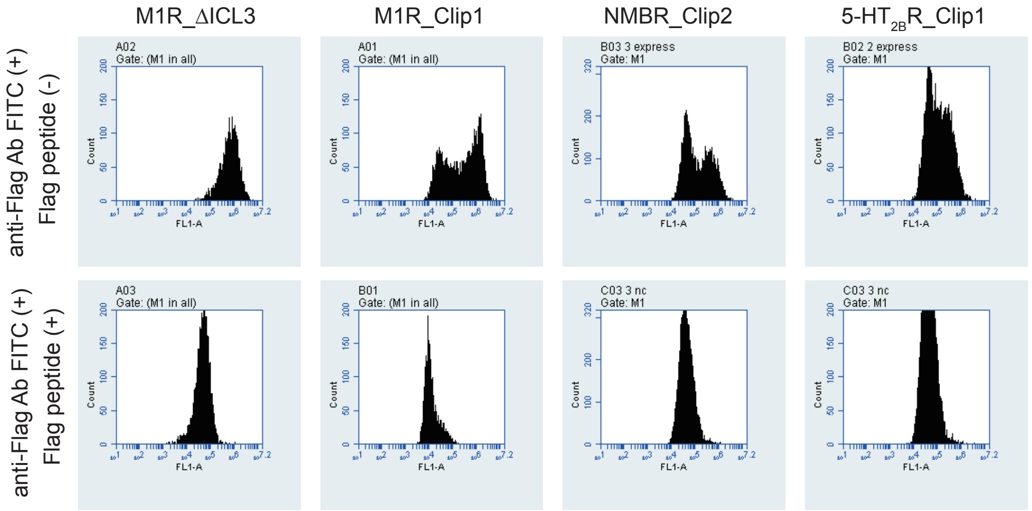


| Mean FITC fluorescence | | | | |
|---|---|---|---|---|
| | M1R_ΔICL3 | M1R_Clip1 | NMBR_Clip2 | 5-HT$_{2B}$R_Clip1 |
| Flag peptide (-) | 941,565 | 691,718 | 333,442 | 240,889 |
| Flag peptide (+) | 61,112 | 18,809 | 69,459 | 57,637 |

**Extended Data Fig. 1 Cell surface stain of different constructs.** The expression levels of different constructs were measured using FITC labelled anti-Flag antibodies (top panels), while the non-specific bindings were measured in presence of 200 μM Flag peptide (bottom panels). The mean FITC fluorescence values are shown in the table.

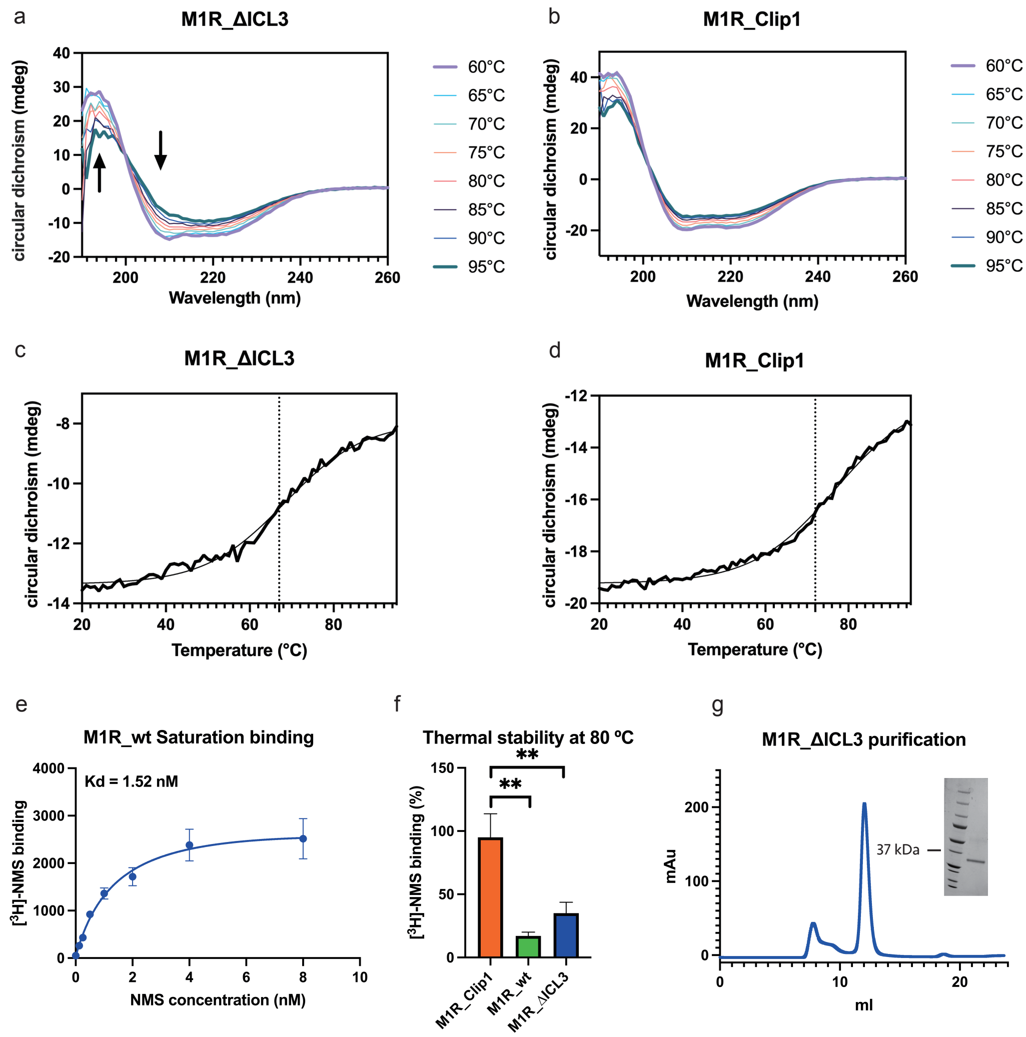


**Extended Data Fig. 2 Biochemical Characterization of the M1R Constructs. a, b** Circular dichroism spectra of the M1R constructs, M1R_ΔICL3 (**a**) and M1R_Clip1 (**b**) at different temperature. **c, d** Protein melting curves at 222 nm. **e** [$^3$H]-NMS Saturation binding assay for the M1R_wt. **f** [$^3$H]-NMS binding assay at 80ºC for 1 min of different M1R constructs in solubilization buffer (blue, M1R_ΔICL3; orange, M1R_Clip1; green, M1R_wt). Values represent the means ± SD of 3 independent experiments. **g** Size exclusion chromatography (SEC) analysis of the M1R_ΔICL3. The peak fraction was analyzed by SDS-PAGE (insert).

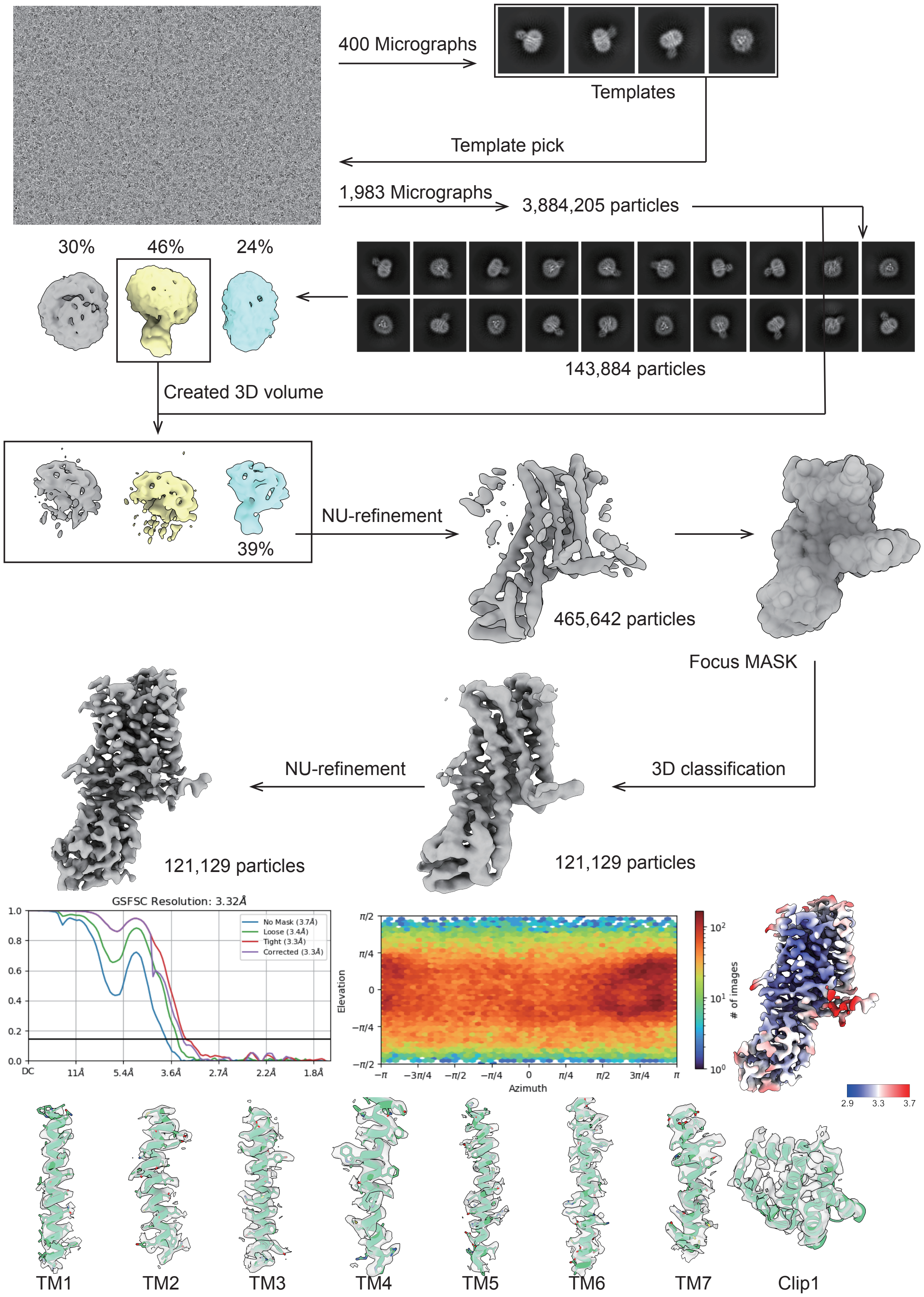


**Extended Data Fig. 3 Data processing of M1R_Clip1.**

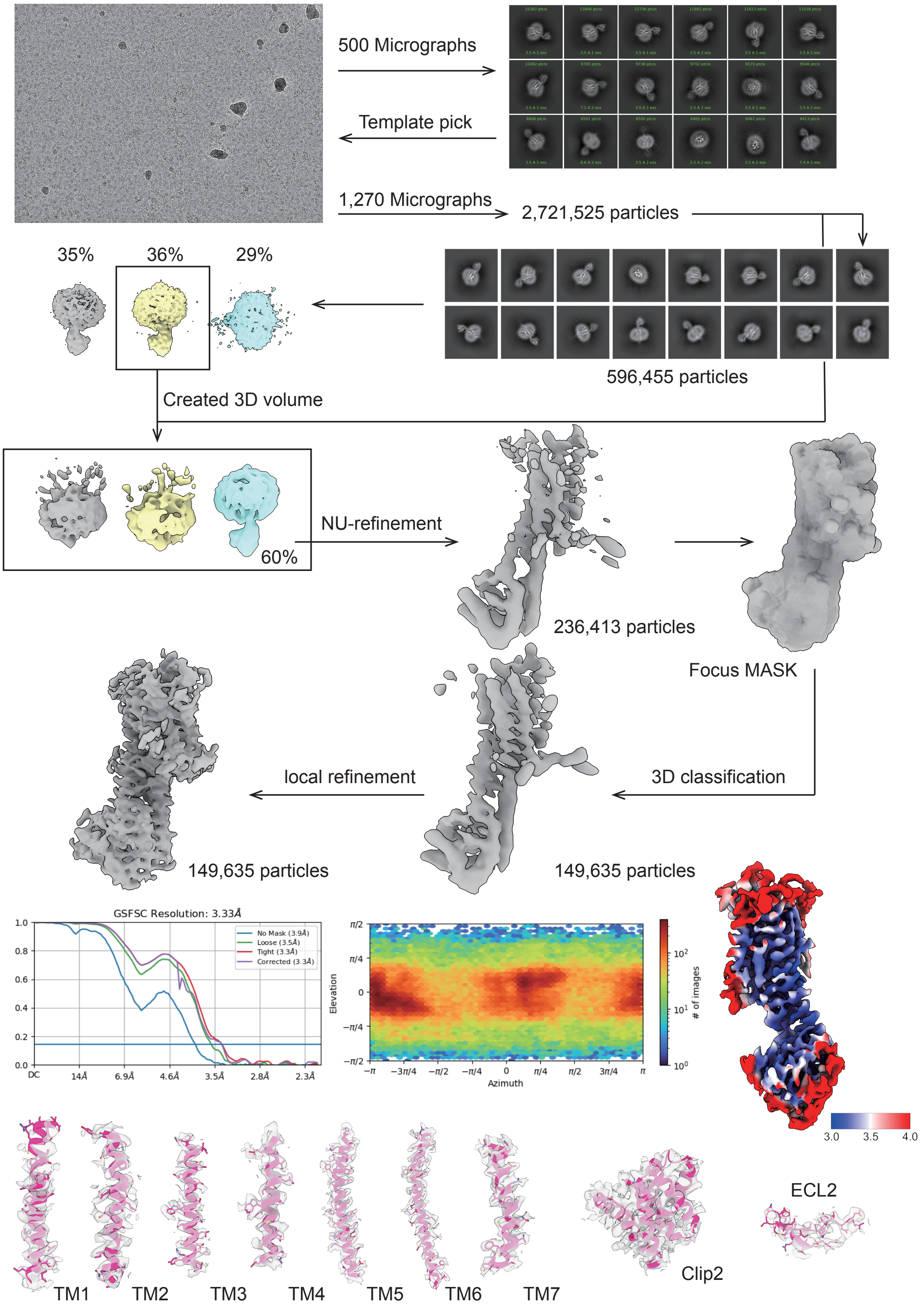


**Extended Data Fig. 4 Data processing of NMBR_Clip2.**

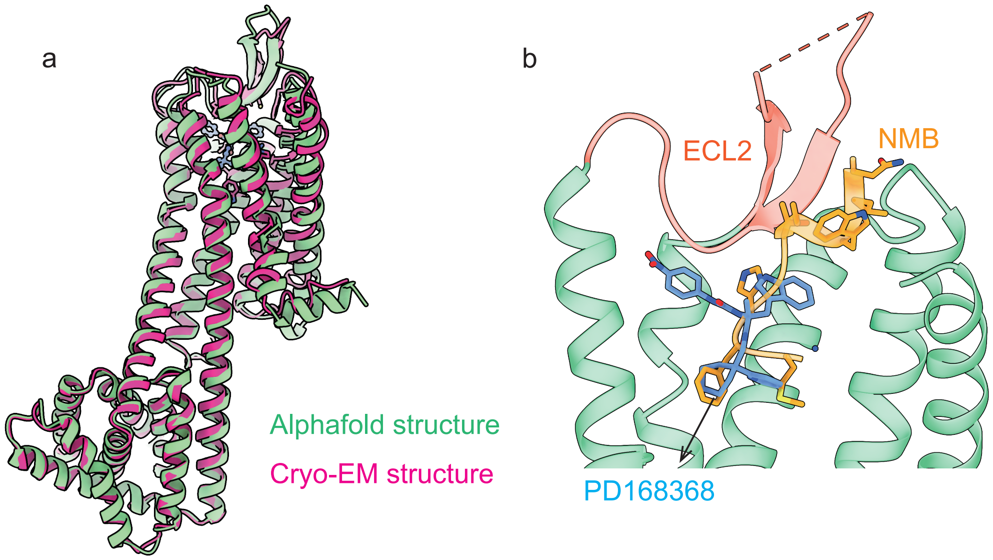


**Extended Data Fig. 5 a** Structure comparison between cryo-EM structure and Alphafold-predicted structure of NMBR_Clip2. **b** Binding pocket comparison between agonist (NMB, colored in orange), and antagonist (PD168368, colored in blue).

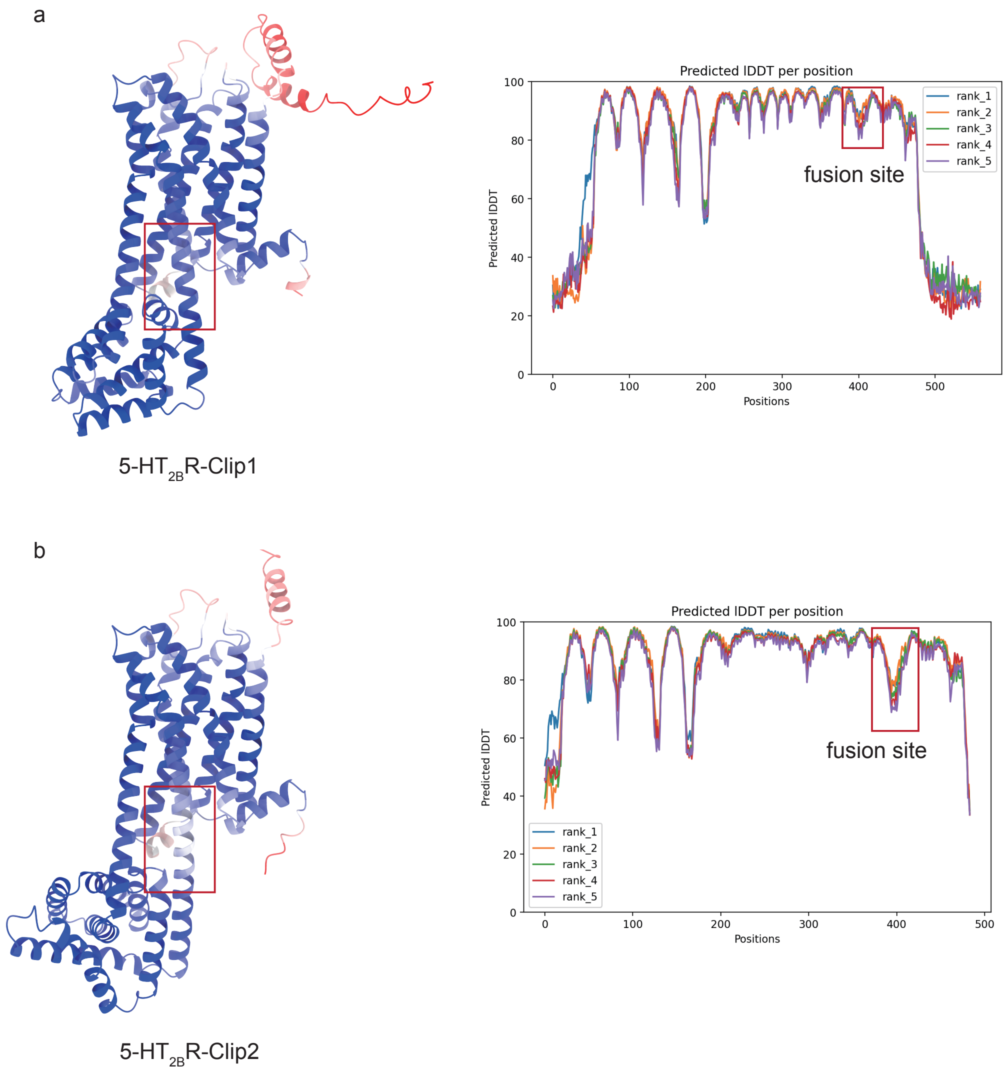


**Extended Data Fig. 6 Representative pLDDT values of Alphafold structure. a** 5-HT$_{2B}$R-Clip1 and **b** 5-HT$_{2B}$R-Clip2. Fusion sites are highlighted with red box. Values represent 5 independent predicted structures.

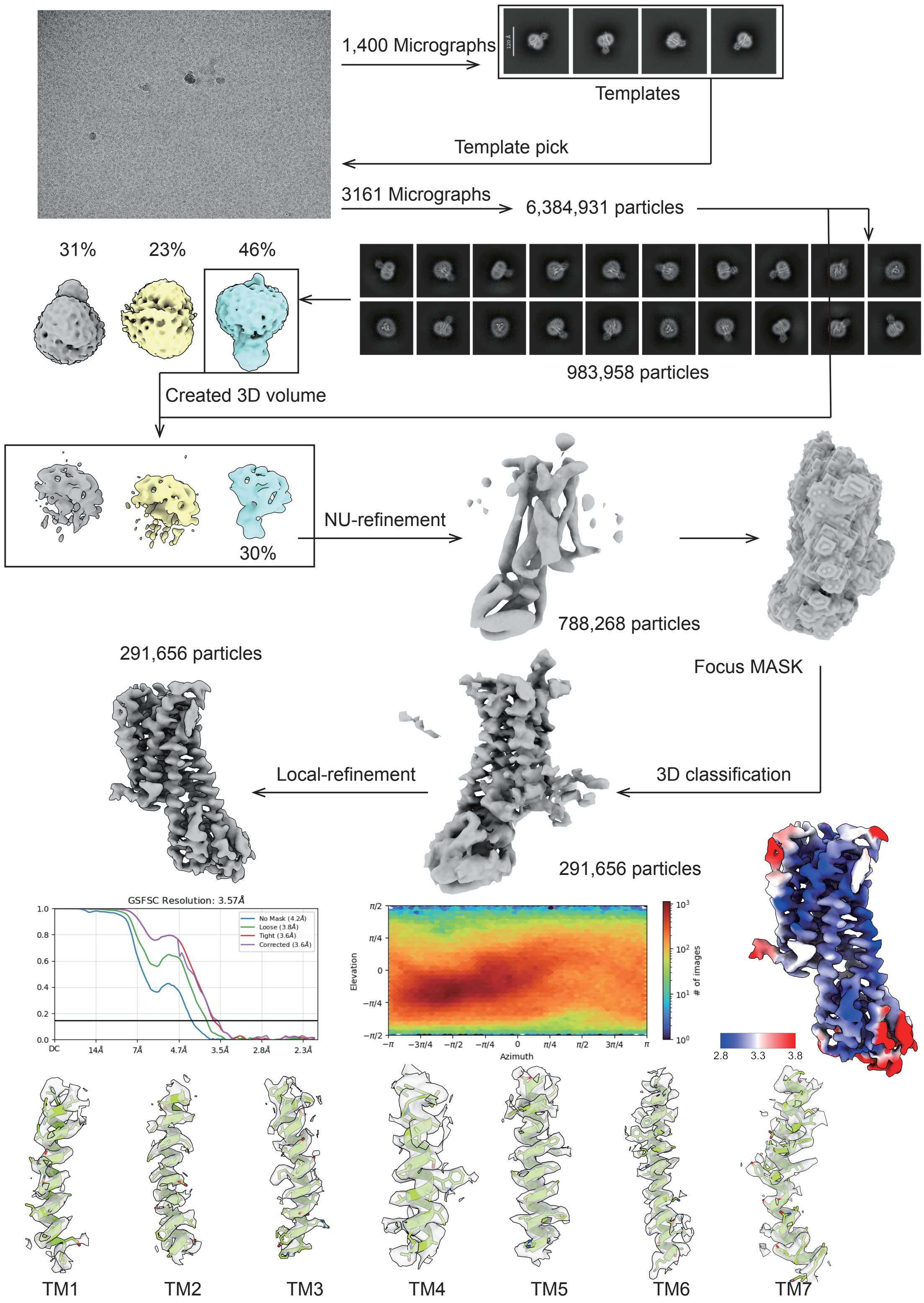


**Extended Data Fig. 7 Data processing of 5-$HT_{2B}$R-Clip1.**
